# Diversity of *Vibrio cholerae* O1 through the human gastrointestinal tract during cholera

**DOI:** 10.1101/2024.02.08.579476

**Authors:** Patrick Lypaczewski, Denise Chac, Chelsea N. Dunmire, Kristine M. Tandoc, Fahima Chowdhury, Ashraful I. Khan, Taufiqur Bhuiyan, Jason B. Harris, Regina C. LaRocque, Stephen B. Calderwood, Edward T. Ryan, Firdausi Qadri, B. Jesse Shapiro, Ana A. Weil

## Abstract

*Vibrio cholerae* O1 causes the diarrheal disease cholera, and the small intestine is the site of active infection. During cholera, cholera toxin is secreted from *V. cholerae* and induces a massive fluid influx into the small intestine, which causes vomiting and diarrhea. Typically, *V. cholerae* genomes are sequenced from bacteria passed in stool, but rarely from vomit, a fluid that may more closely represents the site of active infection. We hypothesized that the *V. cholerae* O1 population bottlenecks along the gastrointestinal tract would result in reduced genetic variation in stool compared to vomit. To test this, we sequenced *V. cholerae* genomes from ten cholera patients with paired vomit and stool samples. Genetic diversity was low in both vomit and stool, consistent with a single infecting population rather than co-infection with divergent *V. cholerae* O1 lineages. The number of single nucleotide variants decreased between vomit and stool in four patients, increased in two, and remained unchanged in four. The number of genes encoded in the *V. cholerae* genome decreased between vomit and stool in eight patients and increased in two. Pangenome analysis of assembled short-read sequencing demonstrated that the toxin-coregulated pilus operon more frequently contained deletions in genomes from vomit compared to stool. However, these deletions were not detected by PCR or long-read sequencing, indicating that interpreting gene presence or absence patterns from short-read data alone may be incomplete. Overall, we found that *V. cholerae* O1 isolated from stool is genetically similar to *V. cholerae* recovered from the upper intestinal tract.

**IMPORTANCE:** *Vibrio cholerae* O1, the bacterium that causes cholera, is ingested in contaminated food or water and then colonizes the upper small intestine and is excreted in stool. Shed *V. cholerae* genomes are usually studied, but *V. cholerae* isolated from vomit may be more representative of where *V. cholerae* colonizes in the upper intestinal epithelium. *V. cholerae* may experience bottlenecks, or large reductions in bacterial population sizes or genetic diversity, as it passes through the gut. Passage through the gut may select for distinct *V. cholerae* mutants that are adapted for survival and gut colonization. We did not find strong evidence for such adaptive mutations, and instead observed that passage through the gut results in modest reductions in *V. cholerae* genetic diversity, and only in some patients. These results fill a gap in our understanding of the *V. cholerae* life cycle, transmission, and evolution.

## INTRODUCTION

*Vibrio cholerae* O1 causes the diarrheal disease cholera, and recent outbreaks are increasing in size and duration^1^. In this context, genomic studies are increasingly conducted to gain an understanding of molecular epidemiology and evolving antimicrobial resistance. Although *V. cholerae* is a small intestinal pathogen, human clinical *V. cholerae* O1 genomes are generated from stool isolates. Gastric acidity kills many ingested *V. cholerae*; however, the proportion that survive can then move into the small bowel^2^. Here, *V. cholerae* can replicate, and the highly motile organisms that locate to small bowel intestinal crypts penetrate the mucin layer overlying the small bowel epithelium and form microcolonies through the action of colonization factors including the Toxin co-regulated pilus (**TCP**)^3^. *V. cholerae* also secrete cholera toxin (CT) that binds to intestinal epithelial cells and stimulates secretion of chloride, causing sodium and water to pass into the intestinal lumen, resulting in diarrhea, vomiting, and dehydration that may be severe. The massive fluid influx into the small intestine can overflow into the stomach and can result in vomiting; the majority of cholera patients experience vomiting during the course of illness^4^ . This is in contrast with other disease processes in which gastric contents alone are vomited. Studies of cholera vomit demonstrate high *V. cholerae* cell counts, and the vomit pH levels approximate the small intestinal environment^5,6^.

*V. cholerae* O1 genomes generated from stool isolates reflect the *V. cholerae* population shed via the large intestine, but *V. cholerae* isolated from vomit may be more representative of the small intestinal *V. cholerae* population that is mediating infection. The genetic diversity of *V. cholerae* O1 could vary along the gastrointestinal (**GI**) tract for several reasons. First, genetic diversity could be reduced through population bottlenecks when a large infecting population is reduced to a smaller number of survivors due to bile and low gastric pH. In animal models, there is an estimated 40-fold reduction in the *V. cholerae* population from the oral inoculum compared to the site of small intestinal colonization^7^. Second, directional natural selection might favor distinct strains from a mixed inoculum. Third, *de novo* mutations, gene transfer events, or gene losses might occur within a patient, some of which might confer adaptation to different niches along the GI tract.

Based on sequencing of *V. cholerae* O1 isolate genomes^8^ and metagenomes^9,10^ from stool, we have previously found that co-infections by distinct strains of *V. cholerae* O1 in humans appear to be rare in the excreted population, but these could be more common in the inoculum, especially in hyperendemic areas. Similarly, the *V. cholerae* O1 population in stool from single individuals contains only minor point mutations and dozens of gene content variants. The level of genetic diversity of *V. cholerae* O1 in the upper GI tract during infection is not known.

To determine the genetic diversity of *V. cholerae* during transit through infected patients, we sequenced *V. cholerae* O1 genomes from paired vomit and stool samples from ten cholera patients in Dhaka, Bangladesh, a region hyperendemic for cholera (**Figure 1**). We compared the single nucleotide variants (**SNVs**) in these 200 genomes and examined variation in gene presence and absence. We show a modest decrease in *V. cholerae* SNVs and gene content variation between vomit and stool, suggesting that passage through the gut does not dramatically reduce genetic diversity or strain variation. We also provide evidence supporting the use of long-read sequencing technologies in accurately determining gene content variation.

**Figure 1.**
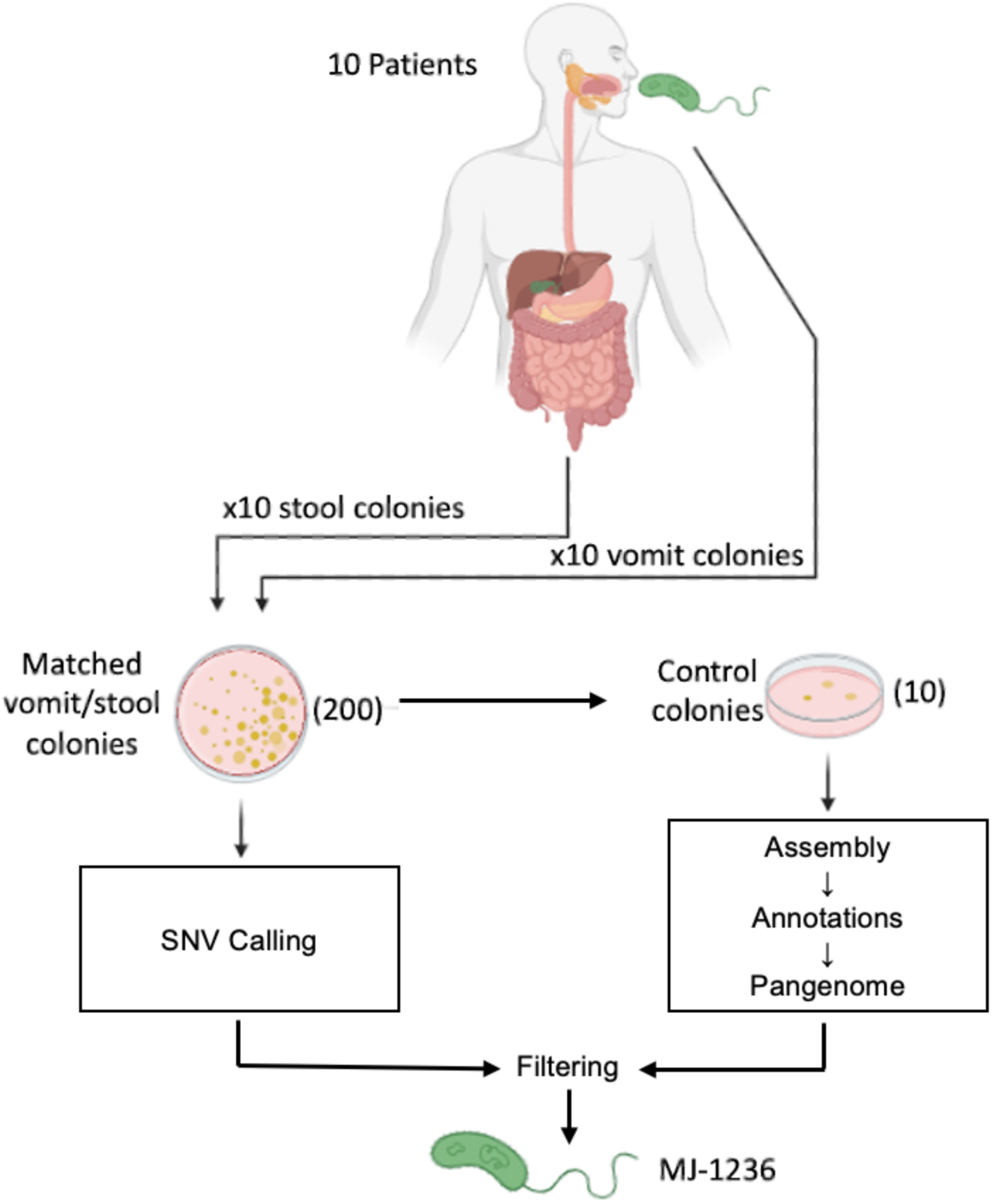
Sampling and sequencing *V. cholerae* O1 isolates from patient vomit and stool. Ten patients were recruited for this study. For each patient, vomit and stool samples were plated to isolate colonies. From each vomit and stool sample, 10 colonies were used for whole genome sequencing in addition to 10 control colonies to evaluate for sequencing errors. Reads were processed independently for SNV calling and pangenome analysis (gene presence/absence) after assembly and annotations, with control colonies used to set filtering thresholds.

## METHODS

### Sample collection and Vibrio cholerae isolation

To examine within-host *V. cholerae* diversity, stool and vomit samples were collected between April 2018 and August 2019 in Dhaka City, Bangladesh from patients aged 2-60 years with severe acute diarrhea and a stool culture positive for *V. cholerae* O1 who had no major comorbid conditions, as in prior studies^8,9^. All samples were collected at the International Centre for Diarrhoeal Disease Research, in Dhaka Bangladesh (**icddr,b**) following the informed consent process. A maximum of 50 mL of vomit and stool were collected immediately upon admission concurrent with clinical interventions, including rehydration and administration of antibiotics. Samples were immediately frozen at −80 °C.

Vomit and diarrheal stool from Bangladesh were stored at the University of Washington at −80 °C and inoculated directly into alkaline peptone water and streaked onto thiosulfate-citrate-bile salts-sucrose agar, a medium selective for *V. cholerae,* or Luria-Bertani (**LB**) agar and tryptic soy agar containing 5% sheep’s blood. Methods used to isolate *V. cholerae* from these vomit and stool samples have been previously described^5^. After incubation at 37 °C for 24 hours, suspect *V. cholerae* colonies were selected and confirmed for the presence of *ctxA* and *V. cholerae* O1 *rfb* gene by PCR^11^ to confirm identification. Twenty individual confirmed *V. cholerae* O1 colonies from each patient (10 from vomit and 10 from stool) were inoculated into LB broth and grown at 37 °C while shaking aerobically overnight. For each colony, 1 mL of broth culture was stored at −80 °C with 20 % glycerol until DNA was extracted.

As a control for sequencing errors and mutations that could occur within culture rather than within patients, we selected one isolate from one cholera patient vomit sample. The glycerol stock of this isolate was streaked onto LB agar and ten colonies were picked for whole genome sequencing. These colonies were used in subsequent analyses as control colonies (**Figure 1**).

### DNA extraction and sequencing

Bacterial glycerol stocks were streaked on LB agar and incubated at 30 °C for 24 hours, and a single colony was picked and grown in 4 mL LB broth with agitation at 37 °C for 18 hours. Genomic DNA was extracted from each of the 200 isolates and 10 control colonies using the Qiagen DNeasy Blood and Tissue kit with RNAse treatment according to manufacturer’s instructions. DNA was then eluted in molecular grade DNase/RNase-free water. Sequencing libraries were prepared with the Lucigen NxSeq AmpFREE kit, pooled and sequenced at the McGill Genome Centre on one lane of Illumina NovaSeq6000 Sprime v1.5 with paired-end 150 bp reads.

### Sequence alignments and SNV analysis

Either the *V. cholerae* O1 strain MJ-1236^12^ or a *de novo* assembly of the genome from the deeply sequenced colony control was used as a reference genome for analysis. To build a phylogeny, reads were processed using Snippy v4.6.0 with default parameters^13^ and the ‘snippy-core’ command was used to generate a core SNV alignment. IQ-Tree v2.2.2.7 was used to infer a maximum likelihood phylogenetic tree from this alignment^14^. The TPM3u+F+I model was chosen by ModelFinder with bootstrap values determined by UFBoot^15^ and rooted on the reference strain for display using iToL^16^. Demultiplexed paired end reads were aligned to the reference genome using the Burrows-Wheeler aligner v0.7.17 with the Maximal Exact Match algorithm^17^. The alignment files were transformed using samtools v1.17^18^ to generate a pileup file. The variant calls were made using VarScan2 v2.4.3^19^. Samples were excluded from SNV analyses if their breadth of coverage was two standard deviations or more below the median.

SNVs within each patient were extracted using bcftools v1.13 with the command ‘bcftools isec - n-[#samples]’. Only SNVs at >90% frequency and read depth >25 were included. These thresholds were established because they resulted in zero SNVs among the control colony genomes. The final list was manually inspected using integrated genome viewer v2.16.0^20^ to remove any false positives resulting from poor mapping or instances in which one colony per group failed to be accurately called and appeared as intra-sample variation. In samples with less than ten colonies remaining after quality filtering, the number of SNVs was normalized to the number of SNVs per ten colonies (*e.g.* 1/9 becomes 1.1/10).

### Genome assembly and pangenome analysis

Isolated genomes were assembled using Unicycler v0.4.9^21^ in Illumina-only mode. The resulting assemblies were quality controlled using checkM^22^ to estimate genome completeness. Four genomes with a completeness score < 100 (isolates BSC08, GSC06, HVC04 and JSC11) were removed from the pangenome analysis. Gene annotations were performed using Prokka v1.14.5^23^ with the reference MJ-1236 proteome as an additional database using the ‘--proteins’ additional argument to maintain consistent annotations and names. Pangenome determination was performed using Panaroo v1.2.8^24^ using ‘--clean-mode sensitive’ to retain the most genes found. We used the Cochran–Mantel–Haenszel test to measure systematic associations between gene presence/absence in vomit and stool, with each patient treated as an observation and the frequency of each gene’s presence in the vomit and stool derived from the panaroo-generated pangenome table. To identify associations at the level of operons, we used the same implementation of the Cochran– Mantel–Haenszel test and counted observations at the level of operons rather than genes.

### Long-read resequencing

Following analysis using short-read data described above, eight colonies with inferred gene content differences within the *tcp* operon were selected for long-read resequencing using a MinION from Oxford Nanopore Technologies. DNA was re-extracted as above and prepared for sequencing using the Rapid Barcoding Kit (SQK-RBK004) according to manufacturer’s instructions to generate sequencing libraries. The libraries were sequenced on a R9.4.1 MinION flow cell. Raw sequencing data was basecalled and demultiplexed to FASTQ files using guppy v6.3.2 (Oxford Nanopore Technologies) using the model dna_r9.4.1_450bps_sup. Reads were assembled using Flye v2.9.1^25^ and a pangenome analysis performed as described above. The results for the *tcp* operon were manually inspected and compared to the short-read data.

### PCR analysis

Colony PCR was performed on *V. cholerae* O1 DNA extracted by boiling *V. cholerae* O1 isolates in molecular grade water at 95 °C for 10 minutes. Taq 2X Master Mix (New England Biolabs) was used for the reaction and PCR primers are listed in **Supplementary Table 1.** PCR products were run on 1.0% agarose gels along an 1kb ladder. Reference *V. cholerae* O1 strains were used to evaluate the *tcp* operon including PIC018, a clinical strain of *V. cholerae* O1 also isolated in Bangladesh^26^ known to have an intact TCP, and a *tcpA* knockout mutant strain of a *V. cholerae* O1 clinical strain isolated in Haiti^27^ gifted by Brandon Sit and Matthew Waldor.

### Ethics Statement

The Ethical and Research Review Committees of the icddr,b and the Institutional Review Boards of Massachusetts General Hospital and the University of Washington approved the study. All adult subjects provided written informed consent and parents/guardians of children provided written informed consent.

## RESULTS

From each of the ten cholera patients, we isolated ten *V. cholerae* colonies from vomit and ten from stool. All isolates were *V. cholerae* serogroup O1 and serotype Ogawa, ascertained using slide agglutination testing using polyvalent and specific antisera, as in prior studies^28^. Isolates underwent whole genome sequencing for identification of SNVs and gene content variation (**Figure 1**). We performed a phylogenetic analysis to examine relatedness of the isolates and found clustering by patient independent of sample type (**Figure 2A**). The ten control colonies grouped together, separated by very short branch lengths, indicating high-quality sequencing and low false-positive SNV identification (**Figure 2A**). We observed a temporal signal in the phylogeny, with genomes isolated in 2018 and 2019 separated by a long branch with strong bootstrap support of 100. Several patients clustered together by time (*e.g.* A, B, C, D and G, H, I, J), which may suggest common exposures (**Figure 2A**). Instances in which more than one patient’s isolates grouped together on the tree (*e.g.* patients E and F) were generally not well supported by bootstraps, making it difficult to reject a model with a single colonization event per patient.

**Figure 2.**
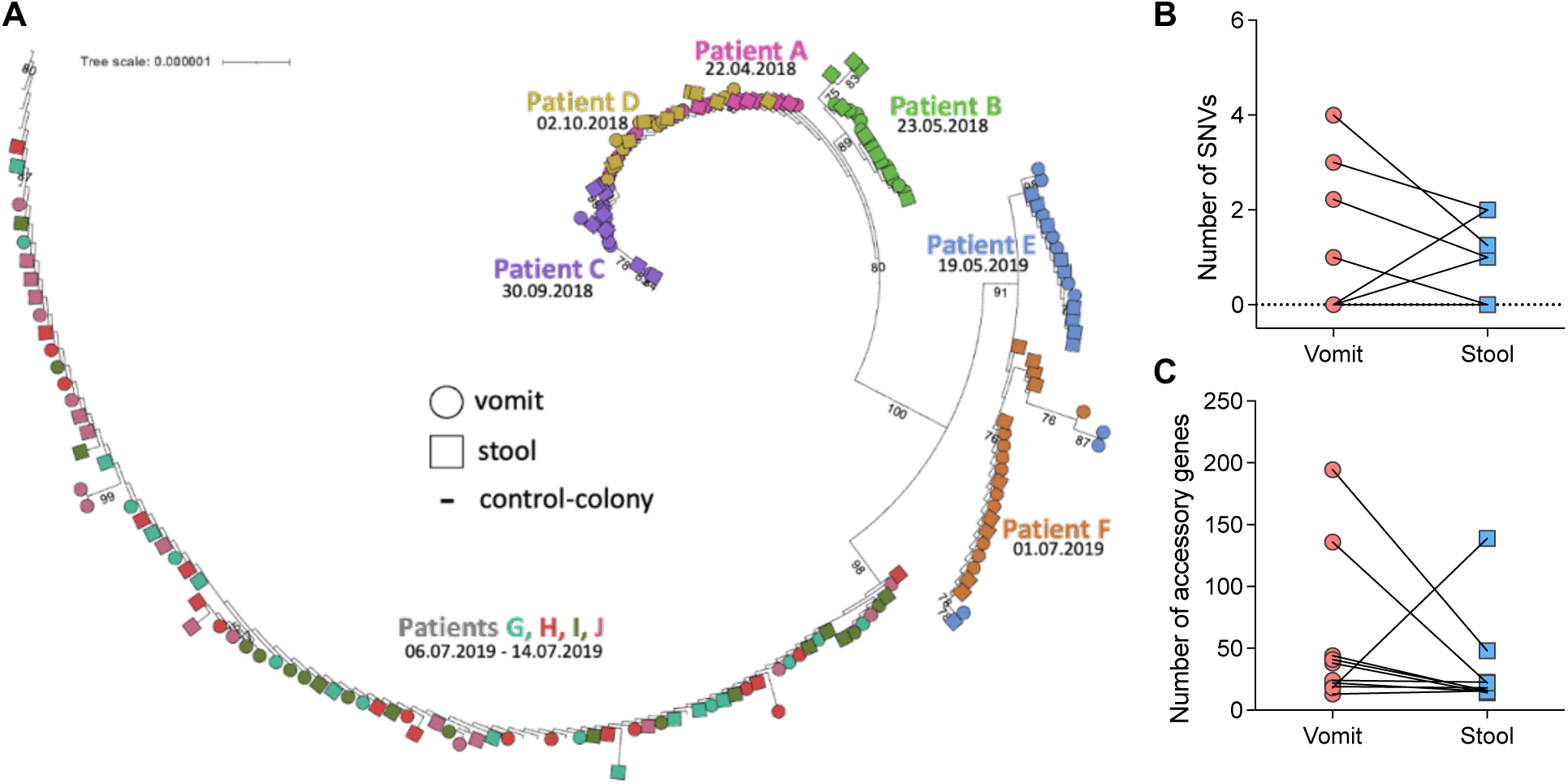
*V. cholerae* O1 within-patient diversity is modestly reduced in stool compared to vomit. **A)** Maximum-likelihood phylogeny of 200 isolates sequenced, demonstrating that isolates cluster by patient and collection date. Ten control colony isolates are also shown. The tree is rooted on the MJ-1236 *V. cholerae* O1 reference genome (branching at the base of patient C). Samples are colored per patient. Collection dates are indicated below patient identifiers. Bootstrap values > 75 are displayed. **B)** Intra-patient variation based on SNVs called against the MJ-1236 reference genome across paired vomit and stool *V. cholerae* genomes, demonstrating a decrease in SNVs in 4 patients, an increase in 3 patients, and no change in 3 patients. The number of SNVs is normalized per 10 colonies, to account for some samples only containing 9 sequenced colonies. **C)** Comparison of within-patient *V. cholerae* gene content in vomit and stool from 10 patients, showing a decrease in 8 patients and an increase in 2 patients. Only ’accessory’ genes that vary in their presence/absence in our dataset are counted here; ’core’ genes common to all genomes are not included.

We next focused on genetic variation using an SNV analysis. Most samples yielded high breadth of coverage (median of 95% to the reference genome) and seven samples with low coverage were excluded (**Supplementary Figure S1**). Because we used multiple media types to isolate *V. cholerae* O1, we tested if the media type was associated with an increase or decrease in intra-sample variation, and found no differences (**Supplementary Table S2**). In comparing vomit and stool within one patient, our findings supported low within-patient diversity. SNVs were always found in a small fraction of colonies from each patient sample (15 SNVs present in 1/10 colonies, 4 SNVs present in 2/10 colonies), and therefore we focused on the number of SNVs per vomit or stool sample rather than their frequencies. If significant bottlenecks occur as *V. cholerae* O1 passed through the gut, we would then expect less genetic variation in stool compared to vomit. Among the seven patients with detectable SNVs, four had fewer SNVs in stool compared to vomit, and three had more (full list of SNVs in **Supplementary Table S3**). Three out of ten patients had no detectable SNVs between vomit and stool isolates (**Figure 2B**). Although the number of SNVs decreased from vomit to stool more often than increased, this difference was not significant (one-sided binomial test, *p* = 0.34). To account for elements of the genome present only in our isolates and not in the MJ-1236 *V. cholerae* O1 reference strain, we repeated an identical analysis using a *de novo* genome assembly using a colony control, which yielded one additional SNV and otherwise identical results (**Supplementary Figure S2**, **Supplementary Table S4**). Thus, we determined that our SNV calling was robust to the choice of reference genome.

In addition to reducing the diversity of point mutations (i.e. SNVs) in a population, bottlenecks would also be expected to reduce pangenome variation. To test this hypothesis, we compared *V. cholerae* gene content variation (e.g., presence or absence) in vomit and stool from the same patient. Based on genomes with high estimated completeness (see Methods), we found a larger total gene content in vomit compared to stool in eight patients, and smaller in two patients (**Figure 2C**; one-sided binomial test, *p* = 0.055). As previously observed^8^, *V. cholerae* O1 gene content is more variable than SNVs within patients, potentially making it easier to detect a change in variation from vomit to stool. Together with the reduction in the number of SNVs from vomit to stool in more patients, these results are consistent with the hypothesis that bottlenecks occur as *V. cholerae* O1 passes through the gut, but that bottlenecks are not evident in all patients and may produce only modest reductions in genetic variation.

We conducted additional analyses to examine the possibility that vomit and stool isolates may experience different selective pressures that select for different genes in the *V. cholerae* O1 population. To identify genes potentially involved in niche adaptation, we tested for genes that varied within patients and were systematically associated with either vomit or stool across patients. We did not find any significant associations at the level of individual genes (Cochran–Mantel–Haenszel test, *p* > 0.05). Because our sample size was likely underpowered to identify gene-specific associations, and the deletion of any member of an operon was likely to disrupt the function of the entire operon, we grouped genes into annotated operons and found that all genes in the *tcp* operon were observed more often in stool than vomit (Cochran–Mantel–Haenszel test, *p* = 0.002). No other significant associations were found. The *tcp* operon encodes the toxin-coregulated pilus, a key bundle forming pilus factor that allows *V. cholerae* to colonize the small intestine^29,30^. Genes essential for human colonization could be stochastically lost from the *V. cholerae* O1 genome in the extra-human environment, and we could expect these loss events would be less common in isolates from stool than the vomit, because the small intestine is a strong selective filter for gut colonization.

To confirm these putative gene loss events, we aligned the raw reads for a set of eight isolates that varied in their presence or absence patterns of *tcp* genes to measure the breadth and depth of coverage of these genes. Genes with substantially reduced breadth of coverage were always called as absent from the pangenome, but the inverse was not always true (**Figure 3A-B**). To further validate these apparent partial deletions, we performed a series of PCRs targeting portions of the genes (primer locations indicated on **Figure 3B**). We were unable to detect any deletion within the *tcp* operon by PCR (**Figure 3C**). To reconcile these conflicting data, we performed long-read resequencing on these eight isolates using Oxford Nanopore Technologies. Interestingly, all eight long-read resequenced isolates contained the entire *tcp* operon, confirming the results of the PCR analysis (**Supplementary Table S5**).

**Figure 3.**
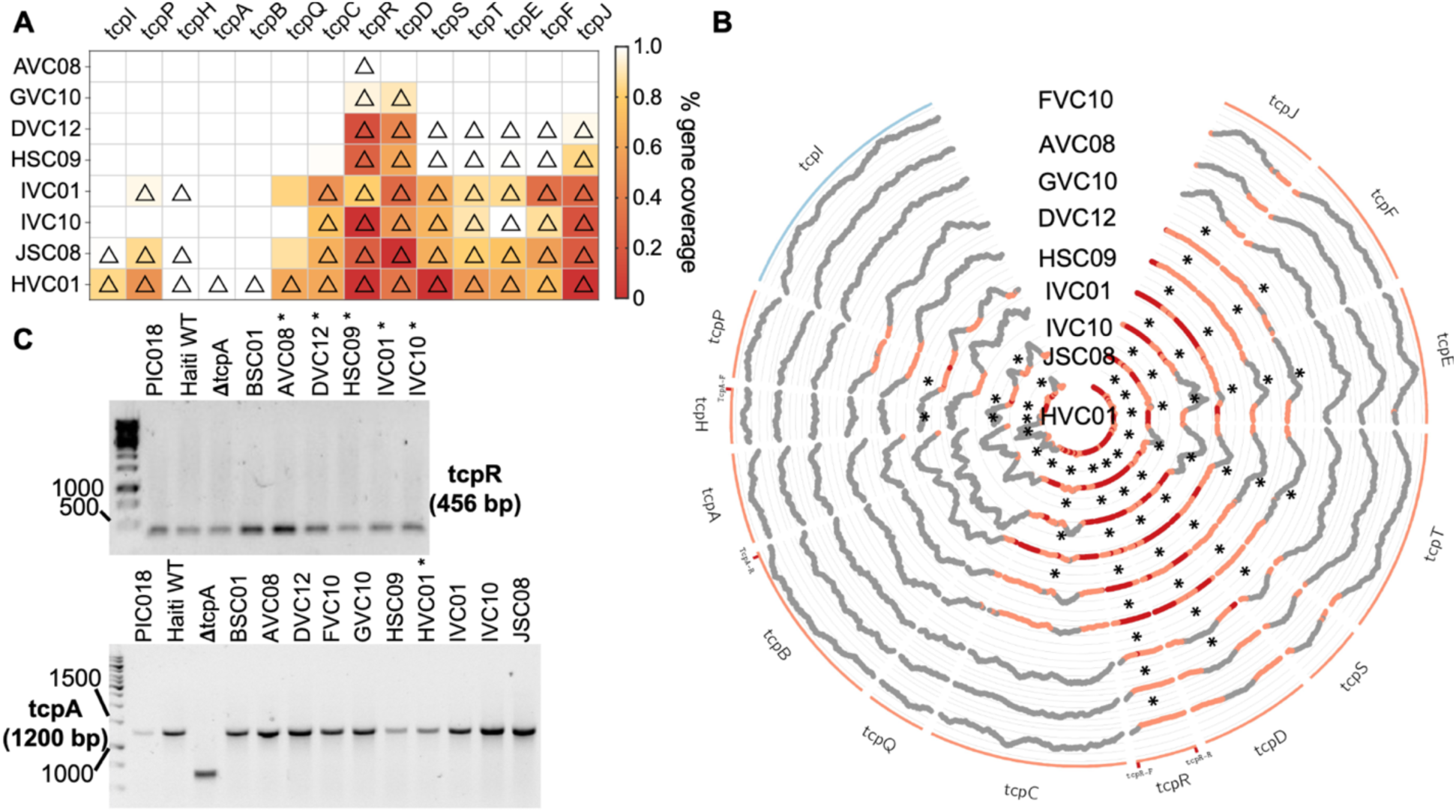
*tcp* genes absent from the *V. cholerae* O1 pangenome have low breadth and depth by short-read coverage and are identified as present by PCR. **A)** The short-read coverage of *tcp* operon genes in 8 genomes with variable presence/absence as determined by panaroo was computed using bedtools and the raw reads aligned to the reference genome *tcp* sequences. Genes called as absent by panaroo are marked with a triangle. The breadth of coverage of each gene is colored according to the scale shown on the right. **B)** Circos plot showing coverage of *tcp* genes. Genes are colored according to the forward (blue) or reverse (red) organization in the genome. The locations of the primers used in PCR assays are indicated on the genes as ‘gene-F/R’. Gene coverage is plotted at each individual base and highlighted in orange at <10% and red at absolute zero. Genes called as absent by panaroo are marked with an asterisk (*). **C)** *tcp* genes assessed in study isolates using PCR. An asterisk (*) indicates expected absence of the gene according to the pangenome analysis. Clinical isolates of *V. cholerae* O1 known to have intact *tcp* and a 𝚫*tcpA* mutant were also used as controls. PCR primers are listed in **Supplementary Table S1.** PCR products were run on 1.0% agarose gels and 1kB ladder.

While these genes often contained frameshift mutations (**Supplementary Table S5**), which might lead to truncated genes, the frameshifts always occurred as part of or immediately downstream of homopolymer sequences that are known to be error-prone in the long-read sequencing^31^.These frameshifts are therefore likely sequencing artifacts. These validation steps suggest caution in interpreting gene presence/absence patterns from short-read data alone.

## DISCUSSION

*V. cholerae* O1 genomic studies in human disease have been based on *V. cholerae* recovered from the stool of patients with cholera, but isolates from vomit may better represent the *V. cholerae* O1 population at the site of active infection in the small intestine. Here, we examined the genetic relatedness between *V. cholerae* O1 recovered from the vomit versus the stool from patients with cholera in Bangladesh. We found an overall low level of genetic diversity between sample types. This suggests that bottlenecks between vomit and stool are not pronounced enough to reduce genetic diversity in the *V. cholerae* O1 population, or that genetic diversity in the initial inoculum is so low that our sample size was insufficient to detect a difference. Additionally, vomiting typically occurs early in the course of infection^4,32^, but does not directly represent the infecting inoculum. Therefore, we cannot exclude a larger bottleneck occurring between the inoculum and the small intestinal *V. cholerae* population. Another explanation for the lack of divergence between *V. cholerae* populations from vomit and stool is that these *V. cholerae* populations are mixed during physiological processes. During high volume fluid secretion by the small intestine, fluid may transit into large intestine and be excreted very rapidly, possibly on the order of minutes to a few hours, effectively homogenizing *V. cholerae* populations across the gut.

Our initial analysis using short-read sequencing suggested that deletions in the *tcp* operon were present more often in vomit than in stool isolates. Because vomit could include ingested environmental strains that may not encode a functional TCP, we thought it was plausible that TCP loss may be observed in vomit isolates more often than in stool isolates^33^. Our results do not fully exclude the possibility that TCP may be sporadically lost by pandemic *V. cholerae* O1 strains in the environment and these genomes would rarely be recovered from humans since they would have impaired ability to colonize the human intestine and survive transit through the GI tract. However, the TCP loss events detected using assembled short reads and pangenome analysis were not confirmed by PCR or long-read sequencing of a subset of genomes. The low depth and breadth of short-read coverage in many of the *tcp* genes suggests that this region of the genome may be difficult to sequence and assemble for technical reasons. It is also possible that our results could also represent within-colony variation in these genes. That is, most cells in a colony contain the intact *tcp* operon (as indicated by their detection by PCR and Nanopore sequencing) but a minority could contain deletions, detectable only by deep short-read sequencing. Such fine-scale variation in *tcp* could be a topic for future investigation. For the purposes of our study, we refrain from drawing firm conclusions regarding natural selection acting on TCP within patients, and we urge caution in interpretation of pangenome variation from short-read data alone.

While short-read Illumina sequencing is highly accurate, it seldom allows genomes to be completely assembled. In contrast, long-read sequencing produces reads with a lower individual accuracy, but helps achieve closed genome assemblies with a clearer determination of a gene’s presence or absence. Of note, while none of our sequenced genomes contained *tcp* deletions, they almost all contained frameshift mutations in at least one region of the *tcp* operon. These frameshifts could lead to truncated genes that might be identified as ‘absent’ in the pangenome. However, a more likely explanation is that these frameshifts are due to sequencing errors. The frameshifts we detected always followed homopolymer repeats, which are known to be error prone in Oxford Nanopore sequencing^31^. It is possible that the next generation of more accurate Nanopore flow cells (R10) combined with multiple rounds of genome polishing could resolve this issue in future studies.

In summary, we observed low within-patient diversity in *V. cholerae* O1 recovered from vomit versus stool, consistent with prior studies examining only stool isolates. This indicates that *V. cholerae* O1 genomes isolated from stool are likely to represent the population at the site of active infection. If population bottlenecks occur between the upper and lower GI tract, they do not appear to have a large effect on *V. cholerae* O1 genetic diversity and are not universal across all patients. We did identify a modest reduction in genetic diversity, particularly in pangenome diversity, in stool compared to vomit, consistent with a non-negligible role for bottlenecks, which could be explored in larger cohorts or time-series studies. Finally, we highlight that gene presence/absence observations based on short-read data should be treated with caution and confirmed by long-read sequencing or other complementary methods.

## Data Availability

The sequencing data generated for all 200 isolates and 10 colony controls were deposited in NCBI Genbank under BioProject PRJNA1046223.

## ACKNOWLEDGEMENTS

We thank the patients of the International Centre for Diarrheal Disease Research, Bangladesh (**icddr,b**) where these samples were collected, and the staff at the icddr,b for data entry and sample collection. We thank Matthew Waldor and Brandon Sit for the contribution of strains. The International Centre for Diarrheal Disease Research, Bangladesh (icddr,b) gratefully acknowledges the government of the People’s Republic of Bangladesh and Global Affairs Canada. This work was supported by the National Institutes of Allergy and Infectious Diseases (R01AI106878 to E.T.R. and F.Q., R01AI103055 to J.B.H. and R.L.; R01A1099243 to J.B.H. and F.Q., K08AI123494 to A.A.W., and T32HD007233 to D.C.), Fogarty International Center (D43TW005572 to T.R.B. and K43TW010362 to T.R.B.), the Government of the People’s Republic of Bangladesh (to the icddr,b), Global Affairs Canada (to the icddr,b), the Swedish International Development Cooperation Agency (to the icddr,b), the Canadian Institutes of Health Research (Project Grant to J.S. and postdoctoral fellowship 187858 to P.L.) and the UK Department for International Development (to the icddr,b). We report no conflicts of interest.

