## Supplementary Figures and Tables for "Diversity of *Vibrio cholerae* O1 through the human gastrointestinal tract during cholera"

### SUPPLEMENTAL TABLES and FIGURES

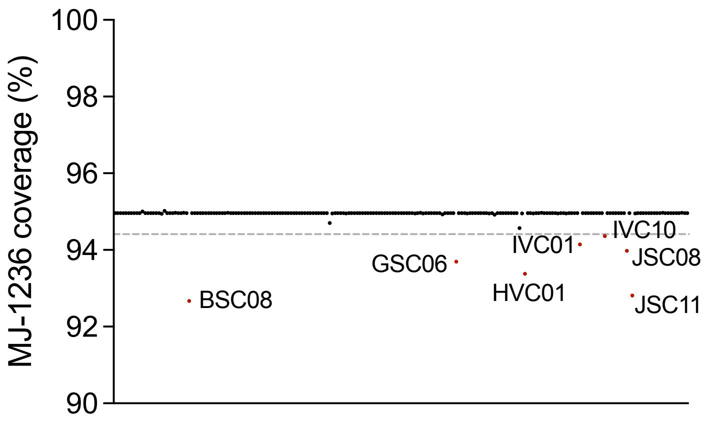


**Supplementary Figure S1. Breadth of coverage of the MJ-1236 *V. cholerae* O1 reference genome.**

The 200 genomes are shown in arbitrary order along the x-axis, with the y-axis demonstrating the breadth of coverage of the MJ-126 reference genome after mapping short reads. Isolates BSC08, GSC06, HVC01, IVC01, IVC10, JSC08 and JSC11 were >2 standard deviations below the median (94.96%) and were therefore excluded from SNV analysis.

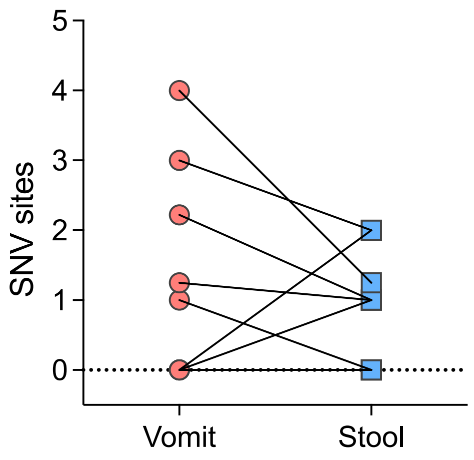


**Supplementary Figure S2. Intra-sample SNVs based on a colony control genome reference.**

Intra-sample variation based on SNVs called against the colony control assembled genome across paired vomit and stool *V. cholerae* O1 populations demonstrate a decrease in diversity in 4 patients, an increase in 2 patients and no change in 4 patients.

**Supplementary Table S1: Primers used for PCR**

| Gene target | Forward Primer (5' - 3') | Reverse Primer (5' - 3') | Expected Amplicon (bp) | Reference |
|---|---|---|---|---|
| *tcpA* | AGCCGCCTAGATAGTCTGTG | TCGCCTCCAATAATCCGAC | 1200 | (1) |
| *tcpR* | CATGACTAGCATATGGTTACATG | TCACATTAACCAAAATACGCC | 456 | (2) |

**Supplementary Table S2. Comparison of media types used to isolate *V. cholerae* O1 from vomit and stool samples and effects on intra-sample SNVs.**

To maximize isolate yield, we used several media types to isolate *V. cholerae* from clinical samples. We found no difference in the number of intra-sample SNVs based on difference in media used for isolation (Chi-squared value = 9.52, df = 10, $p$ = 0.48). **APW** = Alkaline peptone water; **TCBS** = Thiosulfate–citrate–bile salts–sucrose. **TSA** = Tryptic Soy Agar

| Media Type | # of Stool Isolates | # of Vomit Isolates | Isolates with 0 SNVs | Isolates with 1 SNV | Isolates with 2 SNVs |
|---|---|---|---|---|---|
| APW plated onto LB agar | 13 | 18 | 28 | 3 | 0 |
| APW plated onto TCBS agar | 9 | 28 | 33 | 4 | 0 |
| LB agar | 28 | 16 | 42 | 2 | 0 |
| LB broth plated onto LB agar | 14 | 14 | 24 | 2 | 2 |
| LB broth plated onto TCBS agar | 11 | 1 | 11 | 1 | 0 |
| TSA + 5% Sheep’s blood agar | 25 | 23 | 42 | 5 | 1 |

**Supplementary Table S3. SNVs called based on the MJ-1236 *V. cholerae* O1 genome reference.**

The sample types, genes and the SNV effect are shown. Isolates from patients A, B, and G had no intra-sample SNVs. Gene ID and Product annotations are derived from the MJ-1236 *V. cholerae* O1 reference genome.

| Patient | Sample type | Gene ID | Product | SNV |
|---|---|---|---|---|
| C | Stool | VCD_001105 | IS3 family transposase(pseudo) | |
| D | Stool | VCD_001393 | glycosyltransferase family 9 protein | Arg169fs |
| | Stool | VCD_002237 | flagellar hook-basal body complex protein FliE | Ser69Thr |
| | Vomit | VCD_001374 | GDP-mannose 4,6-dehydratase | Thr127fs |
| | Vomit | VCD_001392 | glycosyltransferase family 2 protein | Cys213Tyr |
| | Vomit | VCD_003563 | ribosome biogenesis GTPase Der | Ala146Val |
| E | Vomit | VCD_001366 | acyl-CoA reductase | His664Tyr |
| F | Stool | VCD_002976 | 7-carboxy-7-deazaguanine synthase QueE | Pro217fs |
| | Stool | VCD_001105 | IS3 family transposase (pseudo) | |
| H | Stool | VCD_000480 | retention module-containing protein | Gly452Asp |
| | Vomit | VCD_001352 | mannose-6-phosphate isomerase, class I | Thr74fs |
| | Vomit | VCD_003198 | ATP-dependent Clp protease ATP-binding subunit ClpA | Ala417fs |
| I | Stool | VCD_000299 | phospho-sugar mutase | Pro461Ser |

| | | | | |
|---|---|---|---|---|
| J | Stool | VCD_000300 | amino acid ABC transporter permease | Ser177fs |
| | Vomit | VCD_001362 | glycosyltransferase family 4 protein | Gly232fs |
| | Vomit | VCD_001362 | glycosyltransferase family 4 protein | Leu301Phe |
| | Vomit | VCD_001366 | acyl-CoA reductase | Ser97Tyr |
| | Vomit | VCD_002072 | riboflavin synthase | Ser200Arg |

**Supplementary Table S4. SNVs based on colony control genome reference.**

The patients, source, genes and the effect of the SNV are listed. Isolates from patients A, B, and G had no intra-sample SNVs. Gene ID are based on Prokka annotations of a colony-control genome. Product annotations are derived from the MJ-1236 *V. cholerae* O1 reference genome. *One SNV found here but not in Table S2 is highlighted.*

| Patient | Source | Gene ID | Product | SNV |
|---|---|---|---|---|
| C | stool | | | T>A (pseudo) |
| D | Stool | CC_02_02790_gene | glycosyltransferase family 9 protein | Arg169fs |
| | | CC_02_01528_gene | flagellar hook-basal body complex protein FliE | Ser78Thr |
| | vomit | CC_02_00670_gene | ribosome biogenesis GTPase Der | Ala146Val |
| | | CC_02_02771_gene | GDP-mannose 4,6-dehydratase | Thr127fs |
| | | CC_02_02789_gene | glycosyltransferase family 2 protein | Cys213Tyr |
| E | vomit | CC_02_02763_gene | acyl-CoA reductase | His664Tyr |
| F | stool | CC_02_00089_gene | 7-carboxy-7-deazaguanine synthase QueE | Pro188fs |
| | | | | TA>T (pseudo) |
| H | stool | CC_02_01149_gene | retention module-containing protein | Gly452Asp |
| | vomit | CC_02_00302_gene | ATP-dependent Clp protease ATP-binding subunit ClpA | Ala417fs |
| | | CC_02_02749_gene | mannose-6-phosphate isomerase, class I | Thr74fs |
| I | Stool | CC_02_00968_gene | phospho-sugar mutase | Pro461Ser |

| | | | | |
|---|---|---|---|---|
| | vomit | CC_02_03009_gene | BREX-1 system phosphatase PglZ type B | Ala272fs |
| J | stool | CC_02_00969_gene | amino acid ABC transporter permease | Ser177fs |
| | vomit | CC_02_02758_gene | glycosyltransferase family 4 protein | Gly192fs |
| | | CC_02_02758_gene | glycosyltransferase family 4 protein | Leu261Phe |
| | | CC_02_02763_gene | acyl-CoA reductase | Ser97Tyr |
| | | CC_02_01672_gene | riboflavin synthase | Ser200Arg |

**Supplementary Table S5. Presence of genes in *tcp* operon of genomes resequenced with Oxford Nanopore Technologies.**

Presence of *V. cholerae* O1 *tcp* operon genes in the 8 resequenced isolates as determined by panaroo. Genes present are labeled with '1'. An asterisk (*) denotes the presence of a frameshift within the gene or 5' of the gene but affecting the start site of the gene. Frameshifts were always associated with a homopolymer sequence.

| Isolate | tcpJ | tcpF | tcpE | tcpT | tcpS | tcpD | tcpR | tcpC | tcpQ | tcpB | tcpA | tcpH | tcpP | tcpI |
|---|---|---|---|---|---|---|---|---|---|---|---|---|---|---|
| AVC08 | 1* | 1 | 1* | 1 | 1 | 1 | 1 | 1 | 1 | 1 | 1 | 1 | 1 | 1 |
| DVC12 | 1* | 1 | 1 | 1 | 1 | 1 | 1 | 1 | 1 | 1 | 1 | 1 | 1 | 1* |
| GVC10 | 1* | 1 | 1* | 1 | 1 | 1 | 1* | 1 | 1 | 1 | 1 | 1 | 1 | 1 |
| HSC09 | 1 | 1 | 1 | 1 | 1 | 1 | 1 | 1 | 1 | 1 | 1 | 1 | 1 | 1* |
| HVC01 | 1 | 1 | 1 | 1 | 1 | 1 | 1* | 1 | 1 | 1 | 1 | 1 | 1 | 1 |
| IVC01 | 1* | 1 | 1 | 1 | 1 | 1* | 1 | 1 | 1 | 1 | 1 | 1 | 1 | 1 |
| IVC10 | 1* | 1 | 1* | 1 | 1 | 1 | 1 | 1 | 1 | 1 | 1* | 1 | 1 | 1 |
| JSC08 | 1* | 1 | 1* | 1 | 1 | 1 | 1* | 1 | 1 | 1 | 1 | 1 | 1 | 1 |
